# Genotype-Specific Root Morphology and Metabolic Traits Shape Bacterial Communities and Tolerance to *Fusarium* Root Rot in Wheat

**DOI:** 10.1101/2025.10.15.682011

**Authors:** Omar Hafidi, Marie Simonin, Florent Magot, Yuka Munakata, Alan Kergunteuil, Romain Larbat, Jérémy Grosjean, Alain Hehn, Mathieu Barret, Sophie Slezack

## Abstract

Plant genotype plays a critical role in shaping root-associated microbial communities and in modulating plant tolerance to soilborne diseases such as *Fusarium* root rot (FRR). In this study, we investigated how four wheat (*Triticum aestivum*) varieties, whose tolerance to FRR differs, influence the composition and structure of bacterial communities in the rhizosphere and root endosphere. We evaluated root traits that may contribute to the genotype-specific assembly of bacterial communities across the four wheat genotypes. The variety Concret exhibited the highest FRR tolerance, whereas Pilier was the most susceptible. Analyses of root morphology revealed significant genotype-dependent differences in root length and volume, which were positively correlated with the abundance of some rhizosphere bacteria affiliated with *Bacillus, Lysobacter,* and *Sphingomonas*. Untargeted metabolomics identified 879 features, with 20 key metabolites distinguishing the wheat genotypes, including alkaloids and benzoate- and benzoxazinoid-derived compounds. Correlation analysis revealed significant relationships between these root metabolites and key bacterial taxa. Our findings demonstrate that wheat genotypes influence the assembly of the root microbiota through genotype-based morphological and metabolic traits, providing valuable insights into traits that modulate the plant microbiome to improve wheat resistance to FRR.

## Introduction

Plant roots interact with a multitude of microorganisms in the surrounding soil (rhizosphere) and within their tissues (endosphere). It is now well established that this microbiota plays a functional role in plant phenotype, influencing nutrition, growth and plant health (Berendsen, Pieterse, and Bakker 2012; Dini-Andreote 2020; Mendes et al. 2011). In particular, some results have suggested that both rhizosphere and endosphere bacteria can act as successive layers of protection against fungal root pathogens (Carrion et al. 2019; Wei et al. 2021). Harnessing beneficial plant-microbiota interactions thus provides new approaches for improving plant tolerance to biotic stresses (Becker et al. 2012; Herms et al. 2022). Several factors, including plant species, genotype, and root compartment (rhizosphere versus endosphere), influence the structure of the root microbiota and the relative abundance of beneficial microorganisms (Brown et al. 2020; Cordovez et al. 2019; Quiza et al. 2023). A deeper understanding of how these factors interact is essential for designing root-associated microbiota with desired functional traits. Among these factors, plant genotype plays a central role. Genotypes differing in their tolerance to pathogens often harbor taxonomically and functionally distinct microbial communities in both the rhizosphere and endosphere (Wagner et al. 2016; Mendes et al. 2018; Cordovez et al. 2021; Wagner, 2021). Plant genotypes can affect root microbial communities through variations in their phenotypical traits, including root morphology (length and thickness) as well as the root metabolome (Pascale et al. 2020; Pérez-Jaramillo et al. 2017; Zai et al. 2021). In particular, root diameter, by influencing both plant nutrient and water uptake and C/N resource allocation to the soil, can consequently affect root microbial communities (Herms et al. 2022). Thus, fine roots differ from thick roots in terms of the diversity of rhizospheric microbial communities, as these roots can release more C from rhizodeposits and also specialized metabolites (e.g., benzoxazinoids or coumarins) that influence the recruitment of rhizosphere microbial communities (Herms et al. 2022; Kudjordjie et al. 2022; Mhlongo et al. 2018; Stassen et al. 2021; Stringlis et al. 2019). More broadly, roots, notably in cereals such as wheat, are highly branched, with different types of roots in a single-root system, differing in their age, morphology, anatomy and metabolism (Nakhforoosh et al. 2014). This can therefore modulate the diversity of microorganisms adhering to the root surface and colonizing internal tissues (van der Heijden and Schlaeppi, 2015). Root systems also create a complex chemical environment within root tissues, with a mixture of primary and specialized compounds providing different niches and playing a major role in the assembly of endosphere microbial communities (Pervaiz et al. 2020; Yu et al. 2018).

In this study, our primary objective was to investigate the effects of wheat (*Triticum aestivum*) genotypes differing in their tolerance to *Fusarium* root rot (FRR) on the diversity, structure, and taxonomical composition of microbial communities in the rhizosphere and endosphere. *Fusarium* species, such as *F. graminearum*, are major necrotrophic pathogens of cereals that can infect plants at various developmental stages and cause root diseases, including damping-off and root rot (Cha et al. 2016). Pathogen infection leads to root symptoms such as browning and necrosis and premature plant death (Beccari et al. 2019). We hypothesized that wheat genotypes exert differential influences on rhizosphere and endosphere microbial communities, with certain microbial taxa being preferentially associated with FRR-tolerant and FRR-susceptible genotypes. Our second objective was to characterize both morphological and metabolic root traits that could explain these specific associations among wheat genotypes. We further hypothesized that specific morphological traits and root-derived specialized secondary metabolites contribute to genotype-dependent variations in microbial community composition.

## Materials and methods

### Plant material and soil

The soil used was taken from a 0-15 cm layer in a field at the “Bouzule” research farm (University of Lorraine, 48.74 N, 6.32 E), which is regularly cultivated with winter wheat. The soil characteristics were 42% clay, 46% silt, and 11% sand; the soil pH (water) was 8, the organic matter content was 5.2%, the total N content was 3.2%, and the P_2_O_5_ concentration was 198 mg/kg. After sampling, the soil was sieved at 5 mm and stored at 4 °C until use.

Four winter wheat (*Triticum aestivum*) varieties, i.e., Oregrain, Mutic, Concret and Pilier, were used. Seeds were kindly provided by the Florimond Desprez company (Lille, France). These varieties were initially chosen on the basis of their tolerance profile to *Fusarium* Head Blight (FHB), with Oregrain being the most tolerant, whereas Mutic and Concret were the most susceptible and Pilier showed moderate tolerance to FHB. Their tolerance profile to FRR was evaluated by inoculating germinated seeds with *F. graminearum* conidia at 10^5^.mL^-1^. After 2 hours of incubation at 60 rpm, the inoculated seeds were sown in a mixture of 70% soil and 30% sand (v/v) in a pot (7 x 7 x 8 cm) with 9 seeds per pot. The plants were grown to the 3-leaf stage under the following conditions: 16 h of daylight at 15 °C and 8 h of night at 10 °C, 70% relative humidity, and a light intensity of 300 µmol s^-1^ m^-2^. After 3 weeks of cultivation, the plants were harvested, and FRR disease symptoms were quantified according to the methods of Beccari et al. (2011) using a five-point scale: 0 (symptomless); 1, slightly necrotic; 2, moderately necrotic; 3, severely necrotic; and 4, completely necrotic.

### Experimental design and growth conditions

Seeds of the four wheat genotypes were surface sterilized with 2.4% bleach for 10 min under sterile conditions and then rinsed in two successive baths of sterile ultrapure water for 2 min.

After germination for 3 days at 20 °C in the dark, 9 seeds were transplanted into a pot (7 x 7 x 8 cm) containing 270 g of a mixture of 70% soil and 30% sand (v/v). The substrate was maintained at 70% of its water-holding capacity by weighing the pots every two days and watering them with tap water. The plants were grown to the 3-leaf stage under the following conditions: 16 h of daylight at 15 °C and 8 h of night at 10 °C, 70% relative humidity, and a light intensity of 300 µmol s^-1^ m^-2^. Ten pots were prepared per genotype. At harvest, 5 pots per treatment were randomly selected for further root morphological trait analyses, while the remaining 5 pots were used for metabarcoding and metabolomic analyses. For this latter batch, plants from each pot were randomly divided into two parts, one for amplicon sequencing analysis and the other for untargeted metabolomic analysis.

The root systems were carefully removed from the pots. After the roots were slightly shaken, the soil adhering to the roots, defined as the rhizospheric soil, was collected and stored at −80 °C until processing. All the roots were collected using tweezers and gently washed under tap water. The roots used for morphological trait analysis were stored at 4 °C before analysis (within 2 days after sampling). The roots used for the metabolomic analyses were immediately frozen in liquid nitrogen and stored at −80 °C until use, whereas the roots used for the metabarcoding analyses were surface sterilized before being frozen at −80 °C. For that purpose, after 2 min of incubation in 70% ethanol, the roots were incubated for 5 min in 1.2% bleach. Finally, the roots were washed 3 times for 1 min in sterile water, and the efficacy of root sterilization was checked by plating 100 µl of the last washing water on 10% TSA medium containing 50 mg/L cycloheximide.

### DNA extraction

Root DNA extraction was performed using the DNeasy Plant Pro Kit (QIAGEN, Germany), whereas for rhizospheric soil DNA extraction, the FAST DNA Spin Kit for soil (MP Biomedicals, Unites States) was used (Zai et al., 2021). The amount and quality of the extracted DNA were measured using a Nanodrop 2000 spectrophotometer (Thermo Fisher Scientific, Unites States).

### Metabarcoding amplicon libraries

The target V5-V7 regions of 16S rRNA were prepared using the Metabiote^®^ v 2.0 procedure (GenoScreen, Lille, France) and were paired-end sequenced (2 × 250 bp) using an Illumina MiSeq instrument. Raw sequences were processed into Amplicons Sequence Variants (ASVs) using the following procedure (Callahan et al. 2016). The primers were removed by GenoScreen using CASAVA v1.0. Trimmed fastq files were processed with DADA2v16.1. The following parameters were employed for sequence filtering and trimming: maxN=0, maxEE=c(1,1), and truncQ=5. Chimeric sequences were removed with the *removeBimeraDenovo* function of DADA2. Taxonomic assignment was determined with a naive Bayesian classifier (Wang et al. 2007) using the SILVA_NR99_V138.1 database. ASVs affiliated with mitochondrial and chloroplastic sequences were removed with PhyloSeq (version 1.42). Rarefaction curves were drawn with vegan v 2.6 (Oksanen et al. 2007). This prokaryotic dataset (bacteria and archaea) was rarefied to 2,000 sequences per sample. For all the samples, the rarefaction plateau was reached. Statistical analyses were performed in R version 4.2.2. The downstream analysis was performed with Phyloseq 1.42 and Vegan 2.6.4. Changes in the alpha diversity between plant compartments and varieties were assessed using the Shannon index. A Bray–Curtis dissimilarity matrix and nonmetric multidimensional scaling (NMDS) were used for ordination. The effects of the wheat genotypes on the composition of the bacterial communities were tested by PERMANOVA (using the *adonis2* function). To reduce the complexity of ASV features, PLS-DA was used to identify the most informative ASVs for classifying the four wheat genotypes (Ruiz-Perez et al. 2020). The selected ASVs were tested using ANOVA followed by post hoc Tukey’s HSD test (P < 0.05) to determine group differences.

### Extraction of root metabolites

Two hundred milligrams of fresh root material was ground in liquid nitrogen. One milliliter of aqueous methanol (50%) was added, as was 50 µL of taxifolin (at a concentration of 2 mg/mL), as an internal control. After 10 min of agitation using a vortex, the samples were incubated for 18 h at room temperature. After centrifugation for 10 min at 16,100 g, 600 µL of the supernatant was removed and stored at 4 °C. The remaining pellet was resuspended in 1 mL of 70% aqueous methanol and macerated for 4 h, and after 10 min of centrifugation at 16,100 g, 600 µL of the supernatant was removed. The supernatants were pooled and vacuum dried. The extracts were then solubilized in 300 µL of 70% aqueous methanol. Finally, after centrifugation for 10 min at 16,100 g, the supernatant was filtered using a 0.22 µm filter (Sartorius), dispensed into vials and stored at −20 °C for further UHPLC-HRMS/MS analyses.

### UHPLC-HRMS/MS analysis of root metabolites

Chromatographic analyses were performed on a Vanquish UHPLC system equipped with a binary pump, an autosampler and a temperature-controlled column. Metabolites contained in the extracts (10 µL) were separated on an XB-C18 Kinetex (150 × 2.1 mm, 2.6 µm) (Phenomenex Inc., Torrance, CA, USA) using a gradient of mobile phase composed of mQ water + 0.1% formic acid (A) and LCIJMS grade methanol + 0.1% formic acid (B) at a flow rate of 200 μL·min−1. The elution program started with 10% B for 2 min, then linearly increased from 10% to 30% B in 8 min, and finally increased to 95% B in 10 min. The column was rinsed for 5 min with 95% B and re-equilibrated to the initial conditions for 4 min prior to the next run. The samples were analyzed randomly.

HRMS1 detection was performed on an Orbitrap IDXTM (Thermo Fisher Scientific, Bremen, Germany) mass spectrometer in positive and negative electrospray ionization (ESI) modes. The capillary voltages were set at +3.5 kV and −2.5 kV for the positive and negative modes, respectively. The source gases were set (in arbitrary units min−1) to 40 (sheath gas), 8 (auxiliary gas) and 1 (sweep gas), and the vaporizer temperature was set to 320 °C. Full-scan MS1 spectra were acquired from 120 to 1200 m/z at a resolution of 60,000. MS2 analysis was performed on the pooled sample vials using the data-dependent acquisition (DDA) mode. For this analysis, the AcquireX data acquisition workflow developed by Thermo Fisher was applied. Briefly, this workflow is composed of two main steps and aims to increase the number of MS2 acquisitions, especially at low intensities. First, an inclusion list is generated after the first injection of the sample. Second, the same sample is reinjected, and the ions that have been fragmented are removed from the inclusion list. The second step is repeated six times to ensure the acquisition of MS2 data for a maximum number of ions.

### Metabolomics data processing

The raw UHPLC files were uploaded to Compound Discoverer 3.3 software (Thermo Fisher Scientific, Bremen, Germany) for metabolomic analysis. Briefly, the workflow includes peak detection, chromatogram alignment and peak grouping into features. Each feature corresponds to a specific *m*/*z* at a given retention time. Compounds were identified through elemental composition prediction and searches in public and in-house databases based on mass/formula and MS fragmentation (including LOTUS (Rutz et al. 2022), Chemspider, mzCloud, Mona, and GNPS (Wang et al. 2016)). The detailed workflow used with Compound Discoverer 3.3 is given in the appendix. A mgf file was exported directly from Compound Discoverer 3.3 and imported into SIRIUS (version 5.8.2) (Dührkop et al. 2019). *In*-*silico* dereplication was performed using SIRIUS, and the compound class of each metabolite was predicted with CANOPUS (Dührkop et al. 2021). The similarity between metabolites was assessed using the cossim function of the CluMSID package, after which a molecular network was generated using the igraph package and refined with the Louvain algorithm (Blondel et al. 2008) from the igraph package. Statistical analyses were performed in R version 4.2.2. The MixOmics package was used to analyze the metabolome of the wheat varieties. First, a Principal Component Analysis (PCA) was performed to explore the structure of the metabolome. The varieties were then classified according to their metabolic profile using PLS-DA analysis. Afterward, using the variable importance in projection (VIP score), the metabolic features associated with genotype effects were selected.

### Root trait analysis

Root traits were analyzed as described previously (Romillac et al. 2019). Briefly, roots were suspended in 1-2 cm of distilled water in a 30 x 40cm tray and then scanned at 400 dpi with an Epson Expression 10000XL scanner (Epson, Japan). Images were analyzed with WinRHIZO Reg v.2005c software (Regent Instruments, Quebec, Canada) according to the methods of Regent and Tennant (Arsenault et al., 1995). The total root length (TRL, cm), average root diameter (RAD, mm), fine root length (FRL, diameter < 0.2 mm) and coarse root length (CRL, diameter > 0.2 mm) were measured. The root dry weight (RDW, mg), root volume (RV, cm^3^), volume of fine roots (VFR, cm^3^) and volume of coarse roots (VCR, cm^3^) were also estimated. The root density (RD, g.cm^-3^) was calculated as the ratio of root dry mass to root volume. The specific root length (SRL, m.g^-1^) was calculated as the ratio of root length to root dry mass. Statistical analyses were performed in R version 4.2.2. The root morphological traits were log transformed, and their significance was assessed by ANOVA followed by post hoc Tukey’s HSD test (P < 0.05).

### Linking the wheat microbiome with functional traits

To investigate the relationships between the morphological root traits and the bacterial communities across the four wheat varieties, Redundancy Analysis (RDA) was performed using the vegan package (version 2.6-4). The analysis revealed root traits that differed significantly among the wheat genotypes and ASVs that were significantly enriched among the four wheat genotypes. ANOVA was used to assess the overall significance of the RDA model and to identify the traits contributing significantly to the model.

To analyze the relationships between root metabolites and ASV differing significantly between wheat varieties, Spearman rank correlations were carried out using the *corr.test* function in base R (Knight 1966). The results were visualized using *corr.plot*. To reduce the number of calculations, we considered only the top 20 metabolites according to their VIP scores >1. Correlations with a Spearman coefficient of ≥0.5 or ≤−0.5 and P < 0.01 were considered significant and were visualized in a correlation matrix.

## Results

### Assessment of FRR tolerance, root biomass and root morphological traits of the four varieties of wheat

An evaluation of the FRR symptoms revealed that the four wheat varieties differed significantly (*P=0.004*) in their tolerance to the disease (Table S1). The Concret variety showed a low level of symptoms (mean disease index of 1.3) and was considered the most tolerant to FRR, whereas the Pilier variety appeared to be the most susceptible (mean disease index of 2.3), even though its susceptibility to FRR remains moderate. Compared with these wheat genotypes, Oregrain and Mutic exhibited intermediate levels of susceptibility to FRR (Table S1).

Although the total root biomass (RDW) did not differ among the four wheat varieties, their root systems significantly varied in terms of total root length (*P = 0.0009*) and root volume (*P = 0.02*). These differences were observed in both the fine and coarse root fractions (Table S1). Interestingly, genotypes with greater tolerance to FRR tended to have coarser roots, whereas the more sensitive ones presented finer root systems. Accordingly, in comparison with Pilier, Concret displayed increased coarse root length and volume, along with reduced fine root length (Table S1).

Principal Component Analysis (PCA, Fig. 1) was performed using root traits and the FRR index to compare the four wheat genotypes. The first two PCs explained almost 76% of the total variability. and significantly differentiated (*P=0.01*) the four wheat genotypes; PC1 accounted for 47.8% of the total variation and broadly separated the wheat genotypes according to their sensitivity to FRR on the basis of root traits, especially total root length and coarse root length. The PC2 axis explained 27% of the total variation and mainly opposed Concret, the most tolerant FRR genotype to Pilier, which was the most sensitive genotype in this study, according to the FRR index and root average diameter (RAD) .

**Fig. 1.**
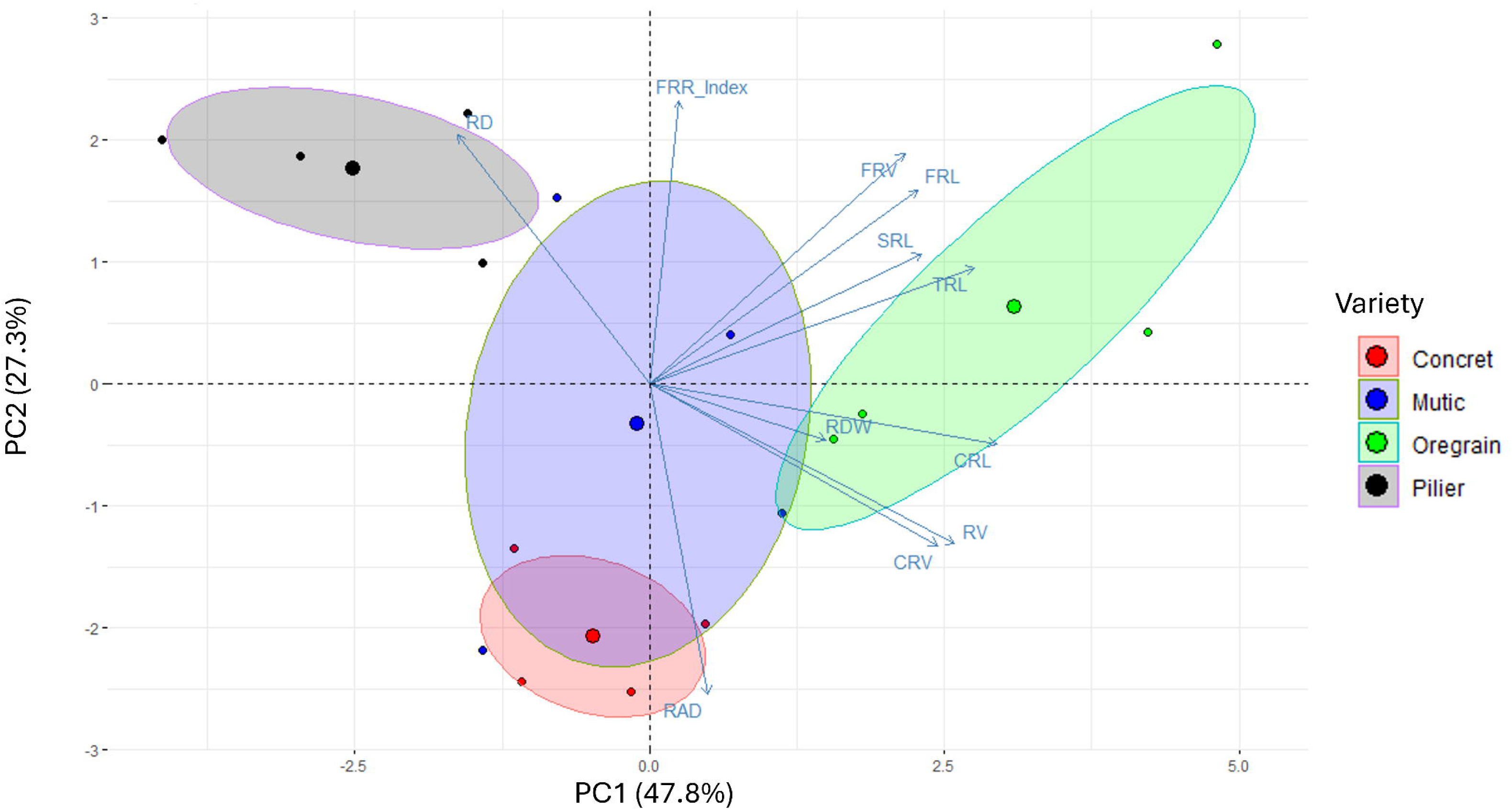
PCA of the effect of root morphological traits and *Fusarium* root rot (FRR) on the four genotypes. TRL: total root length; RAD: root average diameter; RV: root volume; RDW: root dry weight; SRL: specific root length; RD: root density; FRL and FRV the fine root length and volume respectively; CRL and CRV the coarse root length and volume respectively.

### Bacterial diversity in the rhizosphere and the root endosphere of wheat

Given that roots can selectively filter soil microorganisms, we studied the effects of wheat genotypes at an early stage of plant development (3-leaf stage) on bacterial communities, considering both the rhizosphere and root endosphere.

The genotype had no effect on the alpha diversity of either root compartment (Figs. S1 and S2). At the phylum level, the taxonomical composition differed across root compartments. In the rhizosphere, the dominant bacterial phyla included *Bacteroidetes* (22%)*, Actinobacteria* (16.7%), *Proteobacteria* (18.3%), *Gemmatimonadetes* (14%), *Firmicutes* (13.8%), and *Acidobacteria* (7%). In the endosphere, *Proteobacteria* was highly dominant (86%), followed by *Firmicutes* (7%) (Fig. S3).

In the rhizosphere, a significant effect of the wheat genotypes on the structure of the bacterial communities, which explained 25% of the variance, was observed (*P=0.004*; Fig. 2A). A separation between genotypes in terms of almost their sensitivity to FRR was observed on NMDS axis 2. Notably, samples from the Concret and Pilier varieties, which were the least and most susceptible to FRR, respectively, were well separated along NMDS Axis 2. In the endosphere, an even weaker significant effect of the wheat genotypes explained 19% of the variance (*P=0.016*; Fig. 2B). In this root compartment, samples from the variety Concret were separated from samples of the other 3 varieties along NMDS axis 1.

**Fig. 2.**
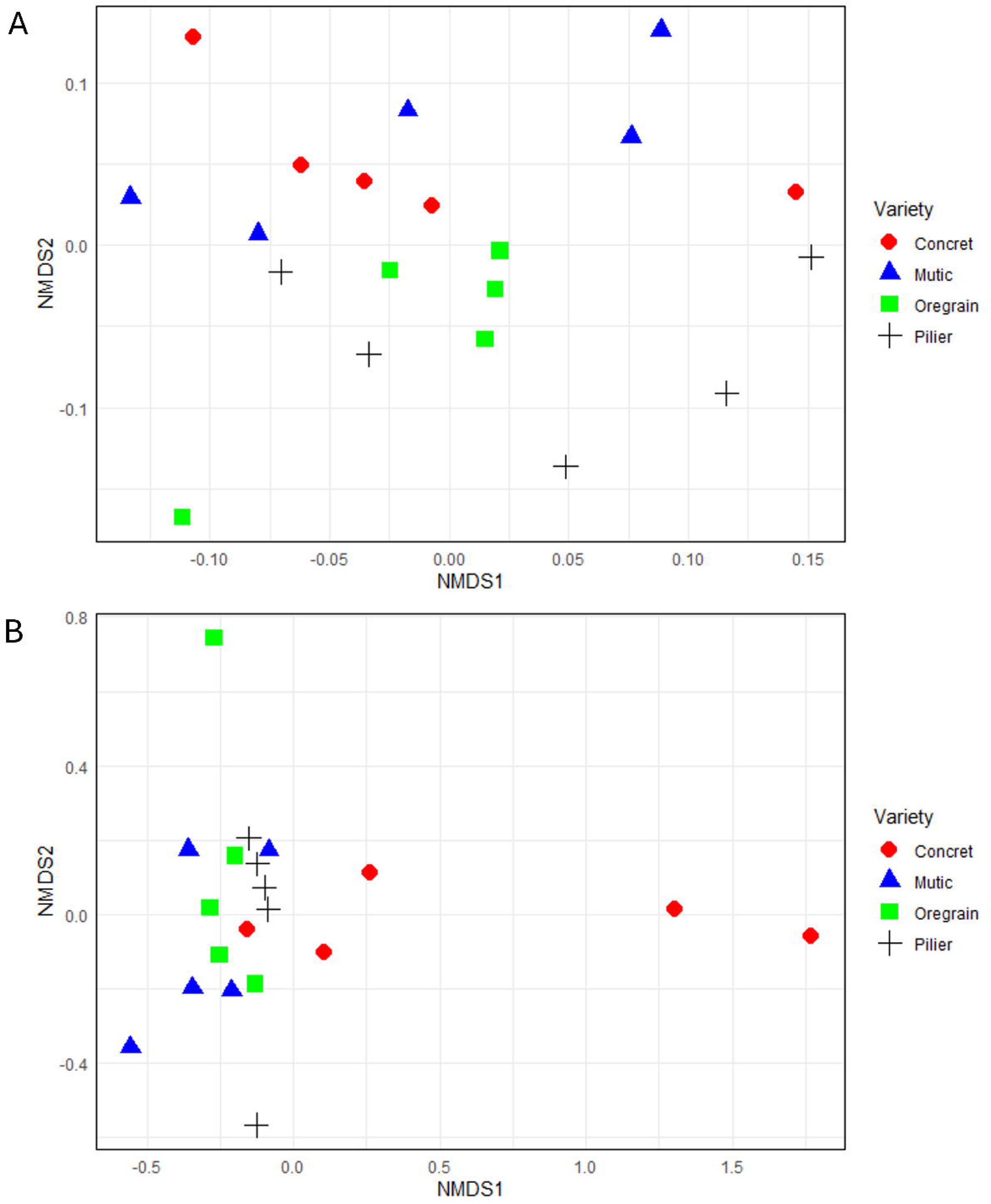
A: NMDS analysis showing the effect of 4 wheat varieties (Concret – Mutic –– Oregrain - Pilier) on the rhizospheric bacterial community structure and **B:** NMDS analysis showing the effect of 4 wheat varieties (Concret – Mutic –– Oregrain - Pilier) on the endospheric bacterial community structure.

We then analyzed which bacterial ASVs were distinct among the four wheat genotypes in each root compartment. In the rhizosphere, 25 ASVs were identified as discriminating, and their relative abundance differed significantly among the four wheat genotypes (Table 1).

**Table 1.**
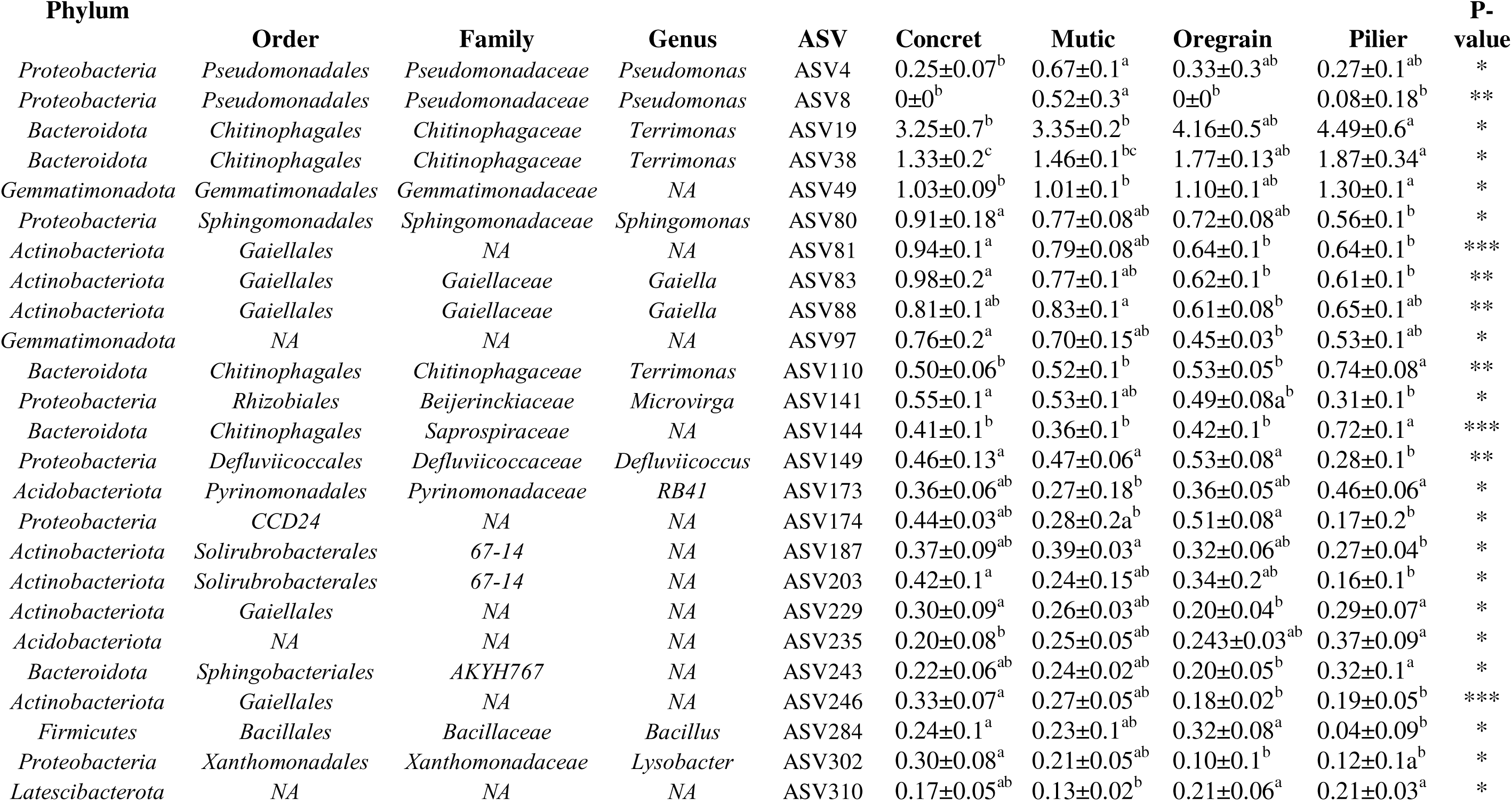
: Bacterial (ASV) with differential relative abundance in the rhizosphere of four wheat varieties (Pv < 0.05). Only discriminative ASVs h mean > 0.05% are displayed. Mean ±standard de viation is shown ( *n* = 5).

**Table 3.**
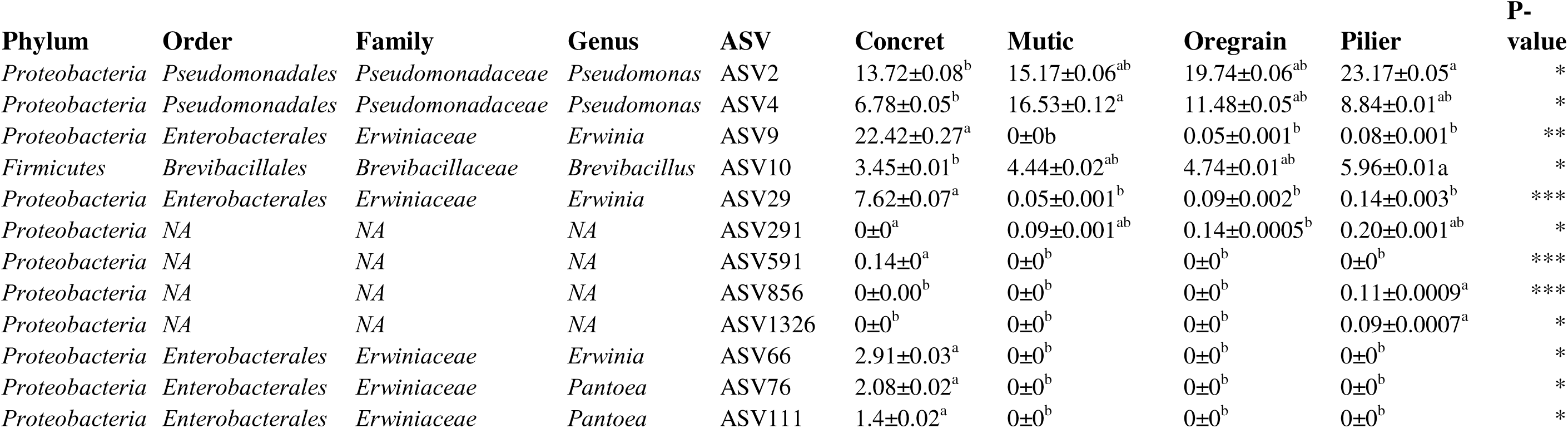
: Bacterial (ASV) with differential relative abundance in the endosphere of four wheat varieties (Pv < 0.05). Only discriminative ASVs h mean > 0.05% are displayed. Mean ±standard de variation is shown ( *n* = 5).

| Phylum | Order | Family | Genus | ASV | Concret | Mutic | Oregrain | Pilier | P-value |
| --- | --- | --- | --- | --- | --- | --- | --- | --- | --- |
| *Proteobacteria* | *Pseudomonadales* | *Pseudomonadaceae* | *Pseudomonas* | ASV2 | 13.72±0.08<sup>b</sup> | 15.17±0.06<sup>ab</sup> | 19.74±0.06<sup>ab</sup> | 23.17±0.05<sup>a</sup> | * |
| *Proteobacteria* | *Pseudomonadales* | *Pseudomonadaceae* | *Pseudomonas* | ASV4 | 6.78±0.05<sup>b</sup> | 16.53±0.12<sup>a</sup> | 11.48±0.05<sup>ab</sup> | 8.84±0.01<sup>ab</sup> | * |
| *Proteobacteria* | *Enterobacterales* | *Erwiniaceae* | *Erwinia* | ASV9 | 22.42±0.27<sup>a</sup> | 0±0<sup>b</sup> | 0.05±0.001<sup>b</sup> | 0.08±0.001<sup>b</sup> | ** |
| *Firmicutes* | *Brevibacillales* | *Brevibacillaceae* | *Brevibacillus* | ASV10 | 3.45±0.01<sup>b</sup> | 4.44±0.02<sup>ab</sup> | 4.74±0.01<sup>ab</sup> | 5.96±0.01<sup>a</sup> | * |
| *Proteobacteria* | *Enterobacterales* | *Erwiniaceae* | *Erwinia* | ASV29 | 7.62±0.07<sup>a</sup> | 0.05±0.001<sup>b</sup> | 0.09±0.002<sup>b</sup> | 0.14±0.003<sup>b</sup> | *** |
| *Proteobacteria* | NA | NA | NA | ASV291 | 0±0<sup>a</sup> | 0.09±0.001<sup>ab</sup> | 0.14±0.0005<sup>b</sup> | 0.20±0.001<sup>ab</sup> | * |
| *Proteobacteria* | NA | NA | NA | ASV591 | 0.14±0<sup>a</sup> | 0±0<sup>b</sup> | 0±0<sup>b</sup> | 0±0<sup>b</sup> | *** |
| *Proteobacteria* | NA | NA | NA | ASV856 | 0±0.00<sup>b</sup> | 0±0<sup>b</sup> | 0±0<sup>b</sup> | 0.11±0.0009<sup>a</sup> | *** |
| *Proteobacteria* | NA | NA | NA | ASV1326 | 0±0<sup>b</sup> | 0±0<sup>b</sup> | 0±0<sup>b</sup> | 0.09±0.0007<sup>a</sup> | * |
| *Proteobacteria* | *Enterobacterales* | *Erwiniaceae* | *Erwinia* | ASV66 | 2.91±0.03<sup>a</sup> | 0±0<sup>b</sup> | 0±0<sup>b</sup> | 0±0<sup>b</sup> | * |
| *Proteobacteria* | *Enterobacterales* | *Erwiniaceae* | *Pantoea* | ASV76 | 2.08±0.02<sup>a</sup> | 0±0<sup>b</sup> | 0±0<sup>b</sup> | 0±0<sup>b</sup> | * |
| *Proteobacteria* | *Enterobacterales* | *Erwiniaceae* | *Pantoea* | ASV111 | 1.4±0.02<sup>a</sup> | 0±0<sup>b</sup> | 0±0<sup>b</sup> | 0±0<sup>b</sup> | * |

Compared with those of other wheat genotypes, the rhizosphere of Pilier was enriched with *Terrimonas* (ASV19; ASV38; ASV110) and a *Saprospiraceae*-related ASV (ASV144). In contrast, *Lysobacter* (ASV302) and *Bacillus* (ASV284) were significantly depleted in the rhizosphere of Pilier. In contrast, Concret, the most tolerant variety to FRR, exhibited higher abundances of *Sphingomonas* (ASV80), *Microvirga* (ASV141) and *Defluviicoccus* (ASV149). The rhizosphere of Concret was also enriched with *Gaiella* (ASV83) and other related Gaiellales taxa (ASV81; ASV229; ASV246).

In the endosphere, Concret was highly enriched in *Erwiniaceae* (ASV9, ASV76 and ASV111). Conversely, compared with other wheat genotypes, the genera *Pseudomonas* (ASV2; ASV4) and *Brevibacillus* (ASV10) were depleted in the endosphere of Concret.

### Comparison of the root metabolomes of the four wheat varieties

A total of 879 metabolic features were detected in root tissues using UHPLC–HRMS. Principal Component Analysis (PCA) was initially performed to compare the root metabolite profiles of the four wheat varieties (Fig. S4). The first two principal components (PCs) explained 30% of the total variance. However, no clear distinction between the wheat varieties was observed. To enhance group separation, Partial Least Squares Discriminant Analysis (PLS-DA) was subsequently performed. The resulting score plot, which explained 21% of the variance, revealed that Concret and Mutic were clearly distinct from each other and from a Pilier/Oregrain cluster (Fig. 3).

**Fig. 3.**
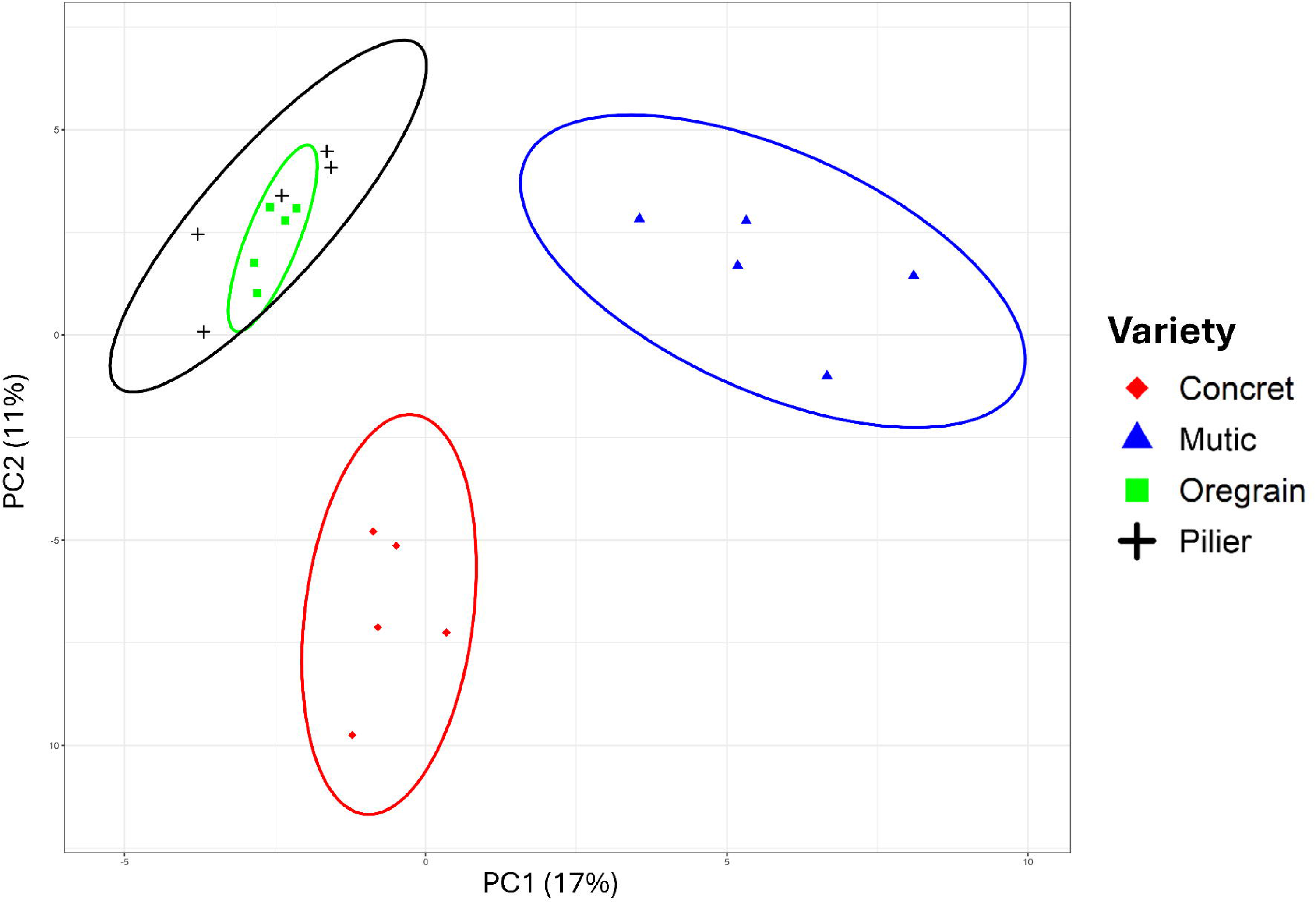
PLS-DA score plot for metabolites in roots of the 4 wheat varieties.

The most discriminant features were identified on the basis of their Variable Importance in Projection (VIP) scores, using a threshold of VIP > 1. Twenty features showed significant differences in abundance among the wheat varieties (Fig. 4) and contributed substantially to genotype discrimination. Although several features remained unidentified, 12 of the 20 could be annotated, at least at the compound family level.

**Fig. 4.**
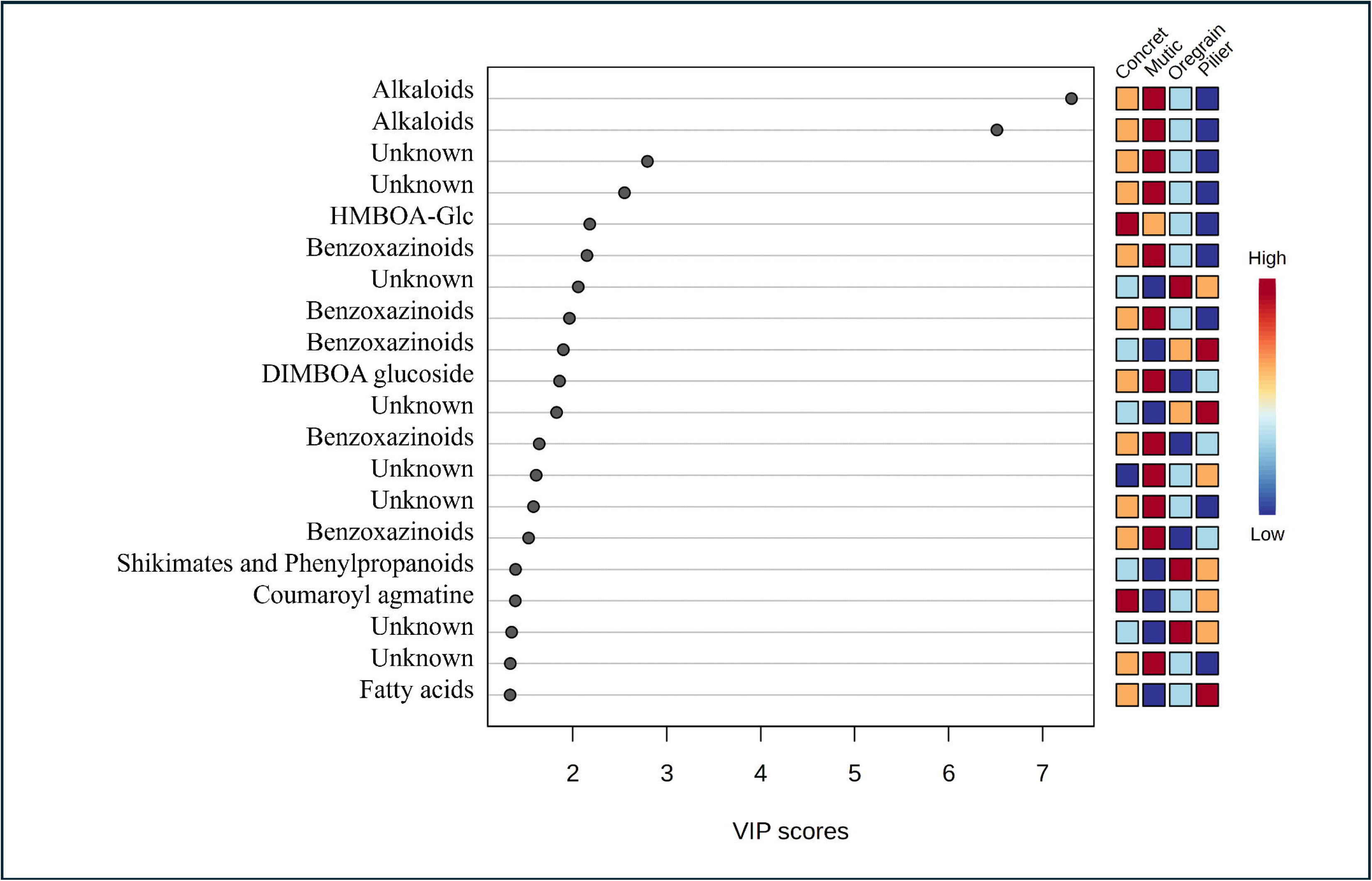
The differential abondance was calculated with variable importance score from PLS-DA. Top 20 metabolites with higher VIP score.

Alkaloids were consistently more abundant in Mutic than in the other varieties, with log2fold changes of 2, 4, and 6.1 relative to Concret, Oregrain, and Pilier, respectively. Moreover, features affiliated with benzoxazinoids were enriched in Mutic and Concret, the most tolerant variety, with an average log2fold change of 1. One signal affiliated with HMBOA-Glc was more abundant in Concret, the most tolerant genotype, while two other signals identified as DIMBOA-glucoside were more enriched in Mutic. Coumaroyl agmatine was more abundant in Concret than in Pilier, with a fold change of 1.5. Unfortunately, the metabolites enriched in Oregrain could not be structurally annotated. However, Shikimates and Phenylpropanoids were more abundant in Oregrain than in the other varieties, with log2fold changes of 0.6 and 2.3, respectively, compared with both Concret and Mutic.

### Linking root morphological traits and bacterial communities

Redundancy Analysis (RDA) was used to assess how root traits influenced discriminating microbial taxa (genus level) among the four wheat varieties. In the rhizosphere, root volume (RV) and fine root length (FRL) were identified as the most influential root traits (*P = 0.009*, R² = 28%, Fig. 5). RV was positively associated with the genera *Bacillus*, *Lysobacter*, and *Sphingomonas* in the variety Concret, the most tolerant genotype, and negatively correlated with Pilier, the most susceptible genotype. Root lengths shorter than 0.2 mm were positively correlated with *Gaiella* and *Terrimonas*, both of which were more abundant in Pilier.

**Fig. 5.**
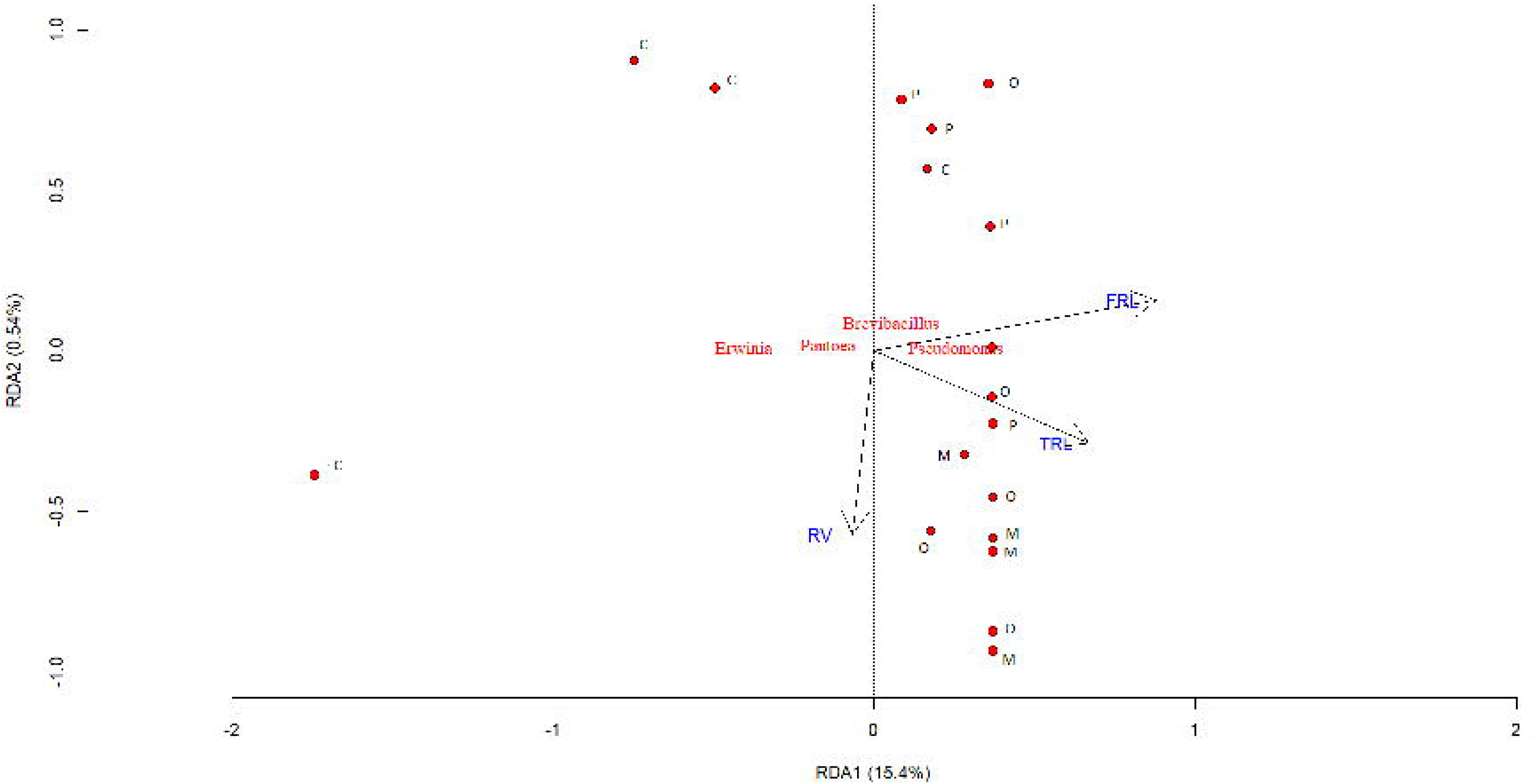
Redundancy analysis of of 16S sequence data and root morphological traits. (a) Root morphological traits as explanatory variables for the divergence between the overall rhizobacterial community composition of the four different wheat varieties (P<0.05). C :Concret, O: Oregrain, M : Mutic and P: Pilier.

**Fig. 6.**
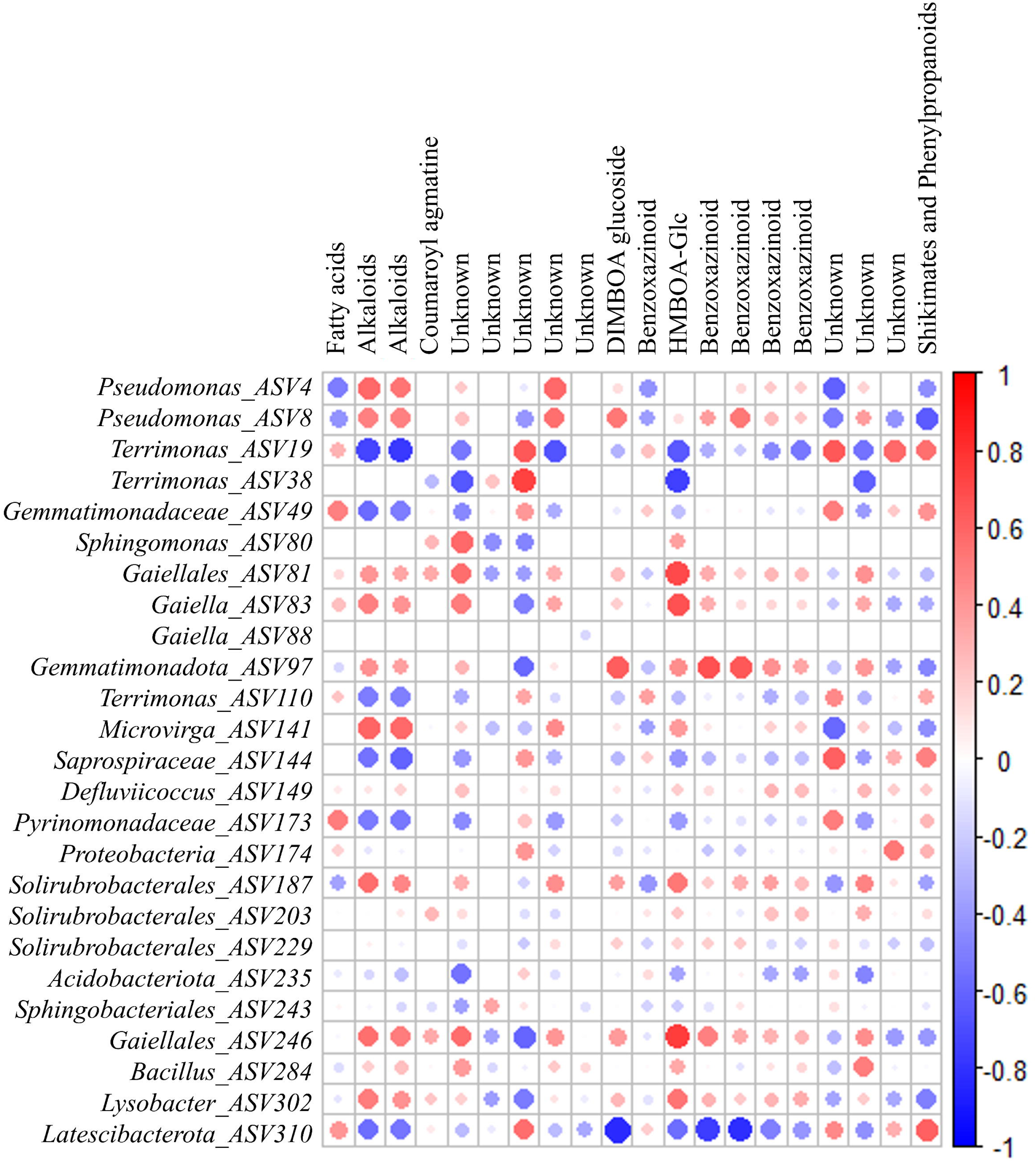
Correlation (Spearman) matrix of rhizosphere bacteria and metabolites in rhizosphere wheat. Only the metabolites that are differentially abundant between the wheat varieties.. High positive correlations are represented by dark blue, negative ones by dark red circles, whereas the circle diameter is indicative for the strength of the correlation. Blank not significantly correlated (< 0.5 or > −0.5 with a *P* > .01).

In contrast, in the endosphere, the RDA biplot revealed that none of the measured root traits significantly explained the variation in bacterial community composition (Fig. S5).

### Correlation between the bacterial community and root metabolites

This analysis included the ASVs that were differentially abundant between the varieties and the top 20 features. Only significant positive and negative correlations with P < 0.01 and |r| > 0.5 were considered (Figs. 7 and 8). In the rhizosphere, a complex network of linear relationships between specific metabolites and bacterial ASVs was observed. Metabolite features related to alkaloids were positively related to *Pseudomonas*-affiliated ASVs (ASV4 and ASV8). Benzoxazinoid-related features were positively related to ASVs affiliated with

**Fig. 7.**
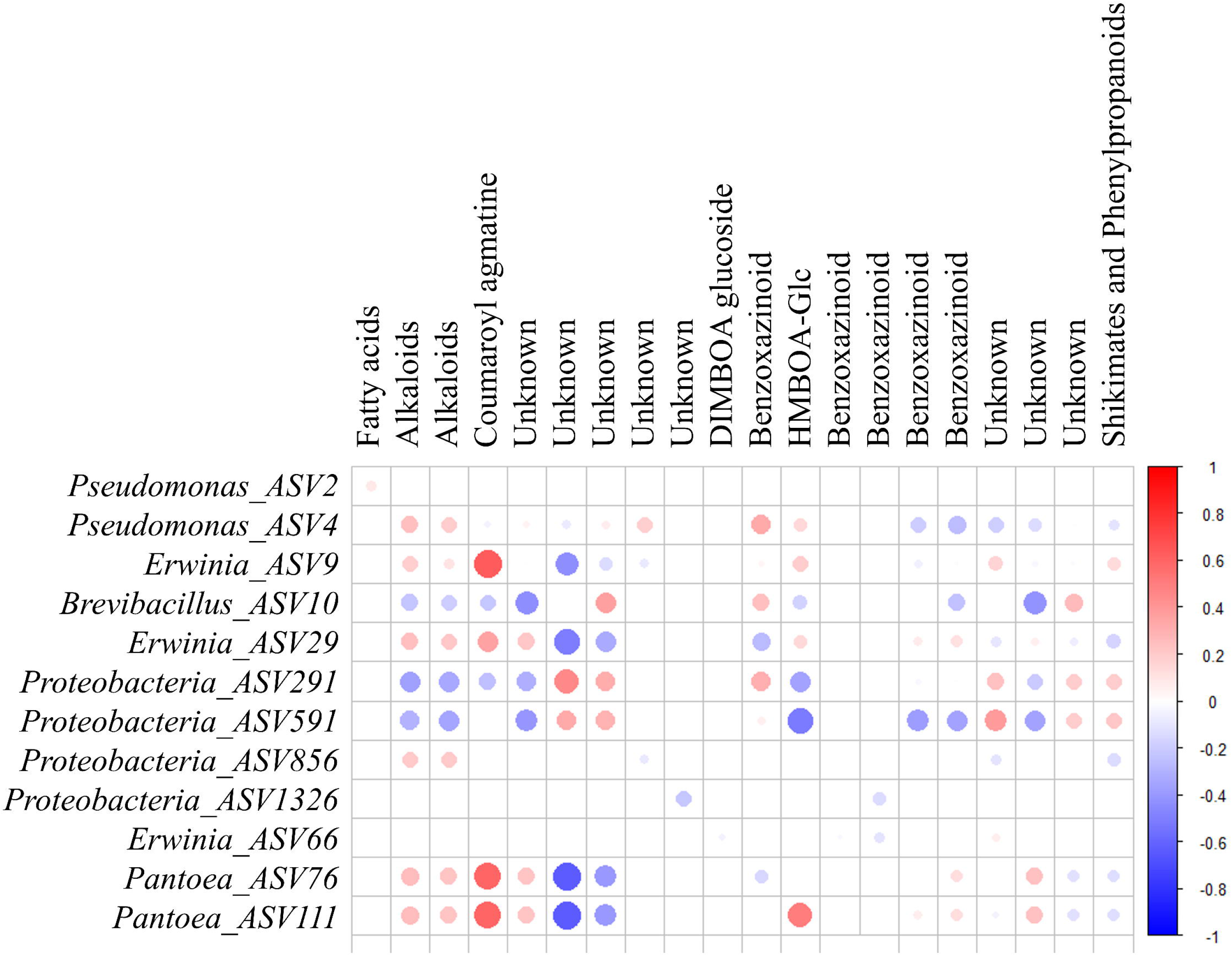
Correlation (Spearman) matrix of endospheric bacteria and metabolites in rhizosphere wheat. Only the metabolites that are differentially abundant between the wheat varieties. High positive correlations are represented by dark blue, negative ones by dark red circles, whereas the circle diameter is indicative for the strength of the correlation. Blank not significantly correlated (< 0.5 or > −0.5 with a *P* > .01).

*Gaiellales*. In particular, DIMBOA-glucoside was strongly positively correlated with *Gaiella* (ASV83 and ASV88) and HMBOA-Glc with ASVs affiliated with *Gaiellales*. In the endosphere, only a few significant correlations were observed. Coumaroyl agmatine was positively correlated with *Erwinia* (ASV9 and ASV66) and *Pantoea* (ASV111).

## Discussion

We compared the rhizosphere and endosphere microbiota of four wheat genotypes whose root traits were characterized and whose tolerance to *Fusarium* root rot (FRR)differed significantly. We attempted to identify specific metabolic and morphological root traits that contribute to the shape of root microbial communities, which could play a role in plant tolerance to FRR.

The clear separation between endosphere and rhizosphere samples confirmed that each plant compartment harbors distinct microbial communities, supporting the view that compartmentalization is a primary factor influencing root-associated microbiota (Bulgarelli et al. 2012; Coleman-Derr et al. 2016; Edwards et al. 2015; Peiffer et al. 2013; Shi et al. 2020).

More notably, our results revealed that wheat genotypes significantly contributed to the structure of the microbial communities within each root compartment (rhizosphere and endosphere) when considered independently. These findings align with those of earlier studies emphasizing the influence of host genotype on the rhizosphere and root microbiota (Brown et al. 2020; Simonin et al. 2020; Quiza et al. 2023). We observed a stronger effect of the wheat genotype on the microbial communities in the rhizosphere than in the endosphere. These findings contrast with those of previous studies reporting more pronounced genotype effects in the endosphere (Quiza et al. 2023; Acuña et al. 2021; Murphy et al. 2021). However, other studies have shown that genotype may also significantly influence community assembly in the rhizosphere (Simonin et al. 2020; Han et al. 2022), suggesting that the strength and direction of genotype effects may depend on other factors, such as plant growth conditions or plant developmental stages (Tkacz et al. 2020; Walters et al. 2018).

Our work also revealed that the clustering pattern of wheat genotypes in the rhizosphere was broadly correlated with their susceptibility to FRR. These findings are in accordance with previous studies showing that plant microbiota can vary in the rhizosphere between disease-tolerant and disease-susceptible plant genotypes (Cordovez et al. 2019; Mendes et al. 2011; Yao & Wu, 2010). The differential recruitment of microorganisms in the rhizosphere could then influence tolerance to FRR. Our results also revealed a similar albeit less pronounced effect in the endosphere. This finding supports the model according to which the rhizosphere and endosphere microbiota function as successive lines of defense against pathogens because of the recruitment of beneficial microorganisms (Carrion et al. 2019).

The composition of the root-associated bacterial communities differed among the four wheat genotypes, particularly in the rhizosphere. In this compartment, a total of 25 ASVs were found to be discriminating among the wheat genotypes. These ASVs, for those identified at the genus level, were affiliated mainly with the genera *Pseudomonas*, *Sphingomonas*, *Terrimonas*, *Microvirga*, *Defluviicoccus*, *Gaiella*, *RB41*, *Bacillus*, and *Lysobacter*. In particular, we observed that the relative abundance of ASVs affiliated with these taxa varied significantly between Pilier, the most sensitive and Concret, the most tolerant wheat genotype to FRR. Pilier harbored a significantly greater abundance of taxa representing the genera *Terrimonas*, *Pseudomonas* and *Brevibacillus*. These genera are well known for their antagonistic effects against plant pathogens (Chandel et al. 2010; Deng et al. 2021; Mehmoud et al. 2022; Ou et al. 2019; Raaijmakers and Mazzola, 2012; Wang et al. 2015). Therefore, the enrichment of the rhizosphere of Pilier, the most sensitive genotype to FRR, among members of these genera raises questions about their functional interactions with *F. graminearum* that remain to be investigated.

In contrast, Concret, the most tolerant genotype to FRR, showed higher abundances of taxa affiliated with *Sphingomonas, Bacillus, Lysobacter and Gaiella*. Most of these genera have been described for their role in pathogen suppression (Asaf et al. 2020; Grady et al., 2016; Hayward et al. 2010; Innerebner et al. 2011; Munakata et al. 2022; Postma et al., 2010; Puopolo et al. 2014). Although the ecological functions of the genus *Gaiella* in plant defense are less well characterized, recent studies suggest that this genus may contribute to soil health and promote plant growth (Alori et al. 2017; Wang et al. 2019). The tolerance of the Concret genotype to FRR could be positively influenced by the relatively high abundance of these specific taxa in the rhizosphere.

In the root endosphere, only a small number of taxa were found to be differentially abundant between wheat genotypes, which is consistent with previous findings that endospheric microbiota are more conserved among genotypes because of stronger host filtering (Edwards et al., 2015). Nevertheless, Concret showed a specific microbial signature, with several ASVs affiliated with *Erwinia* and *Pantoea*, both of which are known to contain members possessing microbial traits to protect plants against pathogens (Duchateau et al. 2024; Gauthier et al. 2019; Guzman et al. 2021; Lopez et al. 2021; Lugtenberg et al. 2016; Smit et al. 2019). Our results suggest a role for these members in the tolerance of Concret to FRR.

The preferential enrichment of specific taxa in the rhizosphere of the wheat genotypes was partly explained by two root morphological traits, i.e., root volume (RV) and fine root length (FRL), as previously observed (Pérez-Jaramillo et al. 2017; Zai et al. 2021). By increasing the amount of soil explored by roots, RV could create heterogeneous physical and chemical conditions in the soil that would promote colonization by different taxa, such as the genera

*Bacillus*, *Lysobacter*, and *Sphingomonas,* in the rhizosphere of Concret. With respect to fine roots, the positive relationship with the abundance of *Terrimonas,* which belongs to Bacteroidetes, in the rhizosphere of Pilier is consistent with previous observations of Pérez-Jaramillo et al. (2017). Compared with those in the rhizosphere, no significant relationships were detected between root morphological traits and specific taxa in the endosphere, suggesting that other factors influence the colonization of internal plant tissues by microbial communities. Finally, the positive relationship between susceptibility to FRR and the length of fine roots suggests that these root morphological traits might impact not only specific taxa but also the pathogen *F. graminearum*. A greater proportion of fine roots, by increasing the availability of nutrients and metabolites, could create conditions that favor colonization by *F. graminearum* (Caldwell 2016; Desgroux et al. 2018).

In addition to variation in their morphology, roots differ in their physiology, and consequently, the composition of root metabolites can vary among different wheat genotypes, influencing the assembly of root-associated microbiota (Pascale et al. 2020). Our results revealed that compared with the most susceptible genotype (Pilier), Concret, the most tolerant variety to FRR, possesses a distinct root metabolome. Notably, higher levels of HMBOA-Glc and coumaroyl agmatine accumulate in the roots of Concret. Hydroxamic acids such as HMBOA and DIMBOA are well known for their antifungal and antimicrobial activities in cereals, contributing to enhanced pathogen resistance (Niemeyer, 2009; Ahmad et al. 2011). Coumaroyl agmatine, a phenolic derivative, is involved in cell wall reinforcement and oxidative defense responses, further strengthening plant defenses (Bassard et al. 2010). These defense-related metabolites could contribute to the better tolerance of Concret to FRR. Furthermore, correlation analyses revealed that HMBOA-Glc and DIMBOA-glucoside were positively associated with b microbial taxa (e.g., *Gaiellales, Erwinia, Pantoea*) enriched in Concret and negatively associated with *Terrimonas* enriched in Pilier, suggesting a role in shaping a more protective microbiota in the rhizosphere of Concret than in that of Pilier.

Benzoxazinoids such as DIMBOA and its derivatives play a central role in modulating plant-associated microbial communities. These compounds act as selective signals and antimicrobial agents, influencing the recruitment and assembly of beneficial microbes in the rhizosphere while suppressing pathogens (Hu et al. 2018; Cotton et al. 2019; Kudjordjie et al. 2019). Seybold et al. (2020) further demonstrated that pathogen-induced shifts in specialized plant metabolism, including benzoxarzinoid pathways, can alter the composition of the plant microbiome.

Interestingly, the root metabolome of Pilier, the most susceptible genotype to FRR, was characterized by relatively high levels of benzoate-related compounds. These metabolites are frequently associated with general plant stress responses rather than active defense mechanisms (Sugiyama, Matsui, & Manabe, 2021). The positive correlation between these metabolites and the relative abundance of *Terrimonas* could be a marker of dysbiosis in the rhizosphere of Pilier.

Collectively, these findings support a scheme in which the increased tolerance of wheat genotypes to FRR could be partly explained by a combination of differences in morphological root traits, the accumulation of root defense-oriented metabolites and the relative abundances of some protective members in the root microbiota.

## Conclusion

Our study highlights the influence of wheat genotypes, which differ in their tolerance to *Fusarium* Root Rot, on the shaping of the rhizosphere and endosphere microbial communities. We observed distinct enrichment of specific taxa in the rhizosphere and endosphere of each wheat genotype. Notably, the most tolerant wheat genotype was associated with specific taxa, such as *Bacillus,* which are well known for their putative beneficial effects on plant health. We also revealed that the recruitment of these specific taxa in the rhizosphere is partly driven by distinct root morphological and metabolic traits that vary among wheat genotypes. In the endosphere, the enrichment of specific taxa appears to be less directly linked to variations in root morphological or metabolic traits. These findings suggest that other factors, such as plant immunity, may play a role in altering the abundance of beneficial microorganisms. Further studies are needed to validate the role of specific microbial taxa and FRR tolerance. Furthermore, our results reinforce the importance of gaining a deeper understanding of plant phenotypic traits that influence the recruitment of specific microbial groups implicated in plant health.

## Author contributions

O.H.: Conceptualization, Formal analysis, Investigation, Methodology, Software, Visualization, Writing – original draft. Y.M : Conceptualization and Methodology. M.S and M.B : Formal analysis, Software of metabarcoding data. F.M, A.K and R.L: Formal analysis, Software of metabolomics data. J.G.: Methodology. A.H.: Funding acquisition, Resources, Supervision, Validation, Writing – review and editing. S.S.: Conceptualization, Data curation, Funding acquisition, Resources, Supervision, Validation, Writing – review and editing. All authors have read and approved the final manuscript version.

## Acknowledgments

We extend our sincere gratitude to PEPLOR for providing access to phytotron facilities, which supported the developmental growth of wheat. We also acknowledge La bouzule center the valuable providing the soil for our experiments.

## Supplementary data

Supplementary data is available at FEMSEC online.

## Conflict of interest

None declared.

## Funding

The authors are very thankful to the financial support provided by the chaire Bio4Solution within the framework of O.H.’s doctoral research project.

## Data availability

**Fig. S5.** Redundancy analysis of of 16S sequence data and root morphological traits. (a) Root morphological traits as explanatory variables for the divergence between the overall rhizobacterial community composition of the four different wheat varieties (P<0.05). C :Concret, O: Oregrain, M : Mutic and P: Pilier.

## Notes

### Competing Interest Statement

The authors have declared no competing interest.

