## Supplementary figures and images for "Genotype-Specific Root Morphology and Metabolic Traits Shape Bacterial Communities and Tolerance to *Fusarium* Root Rot in Wheat"

### Supplementary figure 1.png

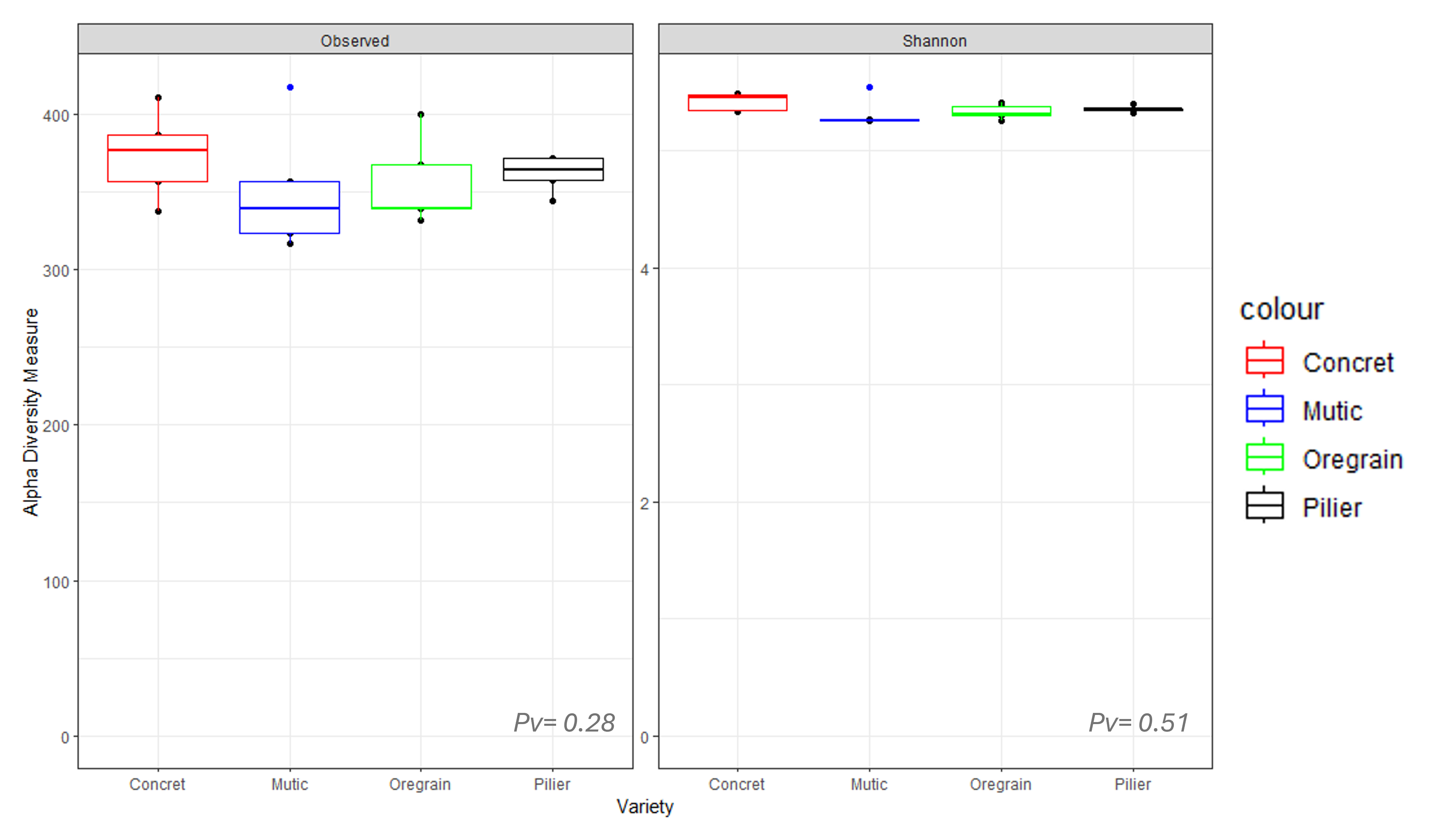

### Supplementary figure 2.png

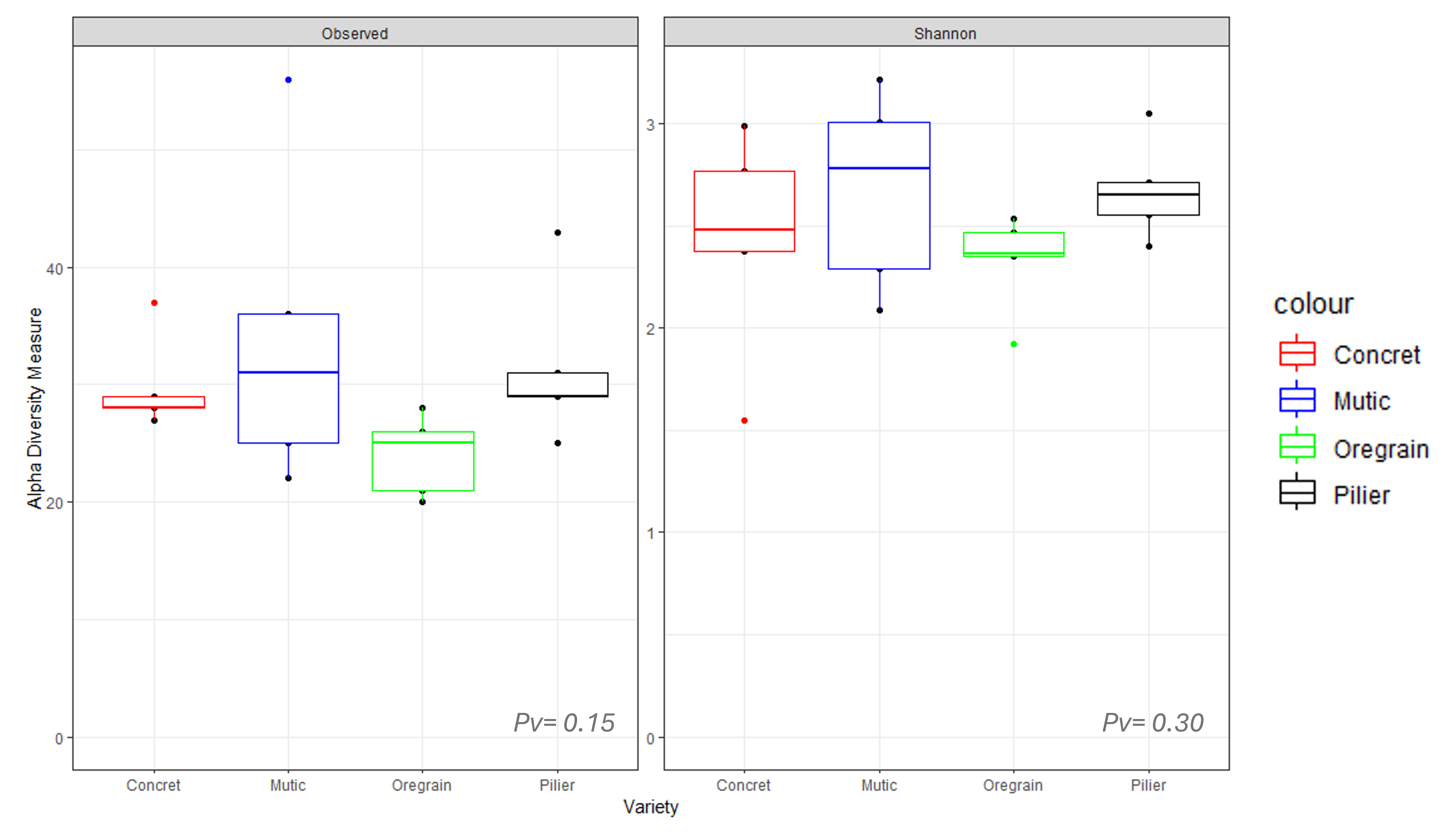

### Supplementary figure 3.png

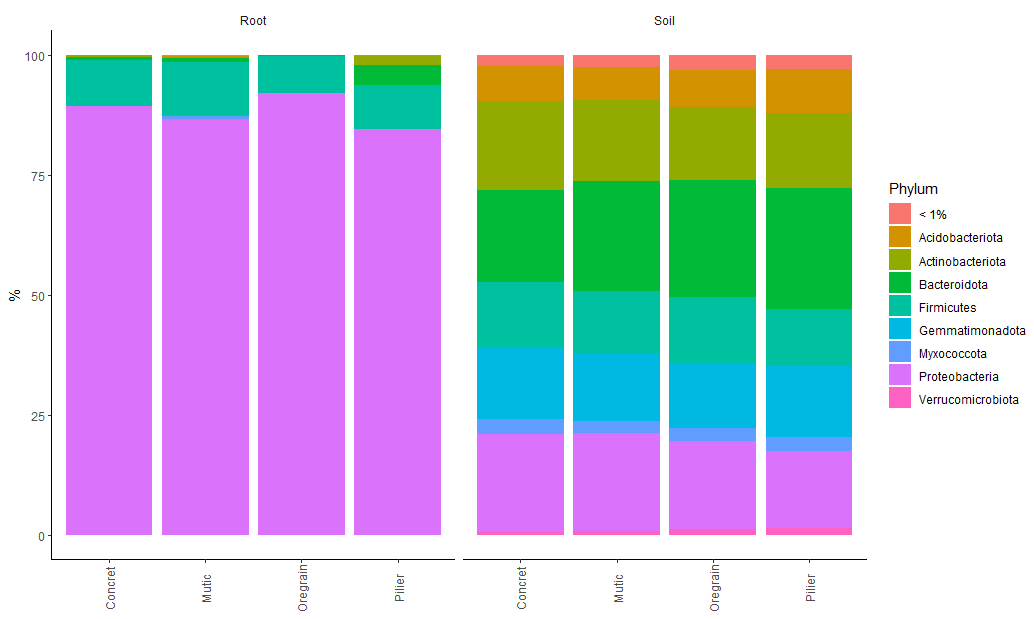

### Supplementary figure 4.png

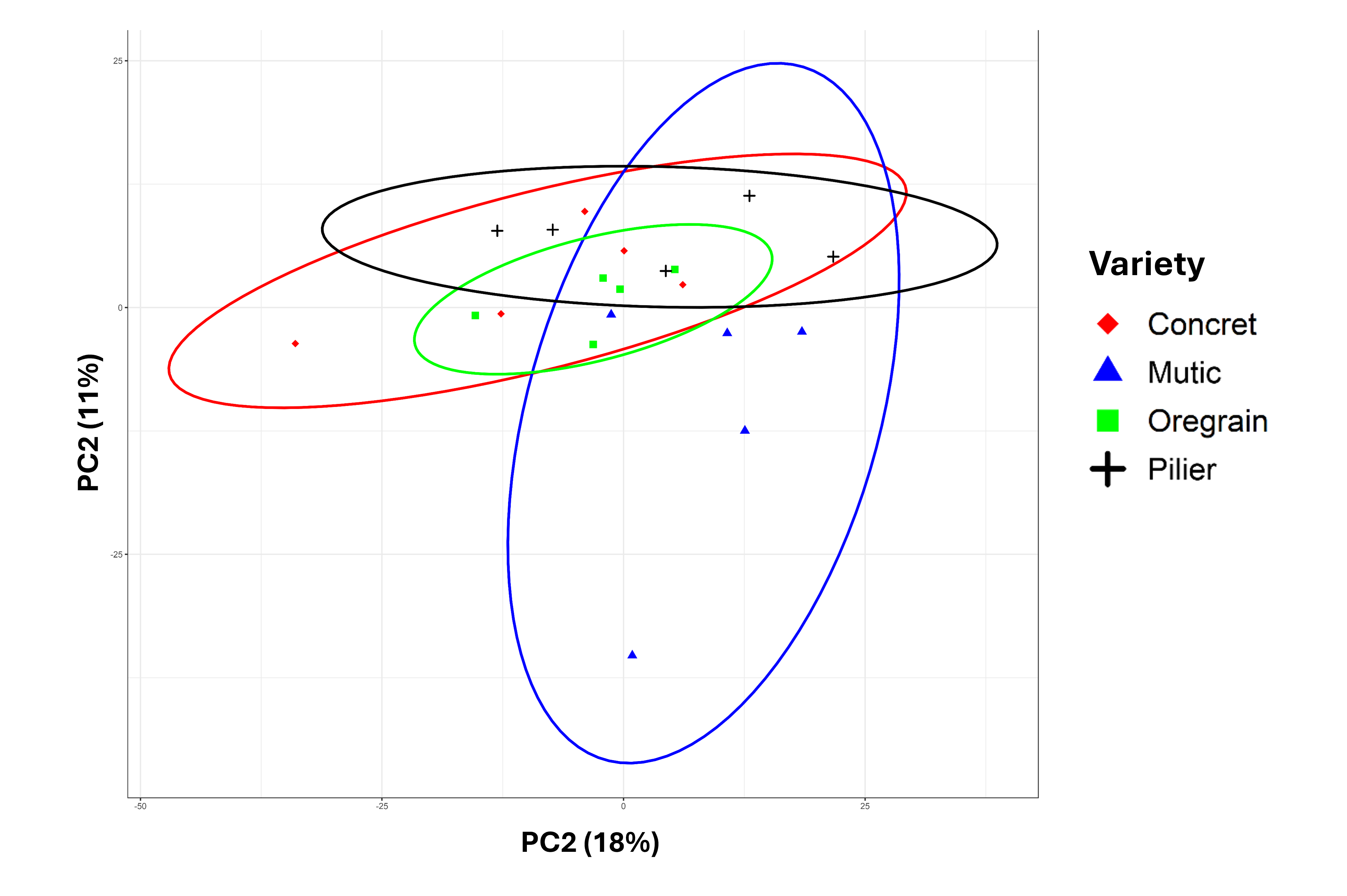

### Supplementary figure 5.png

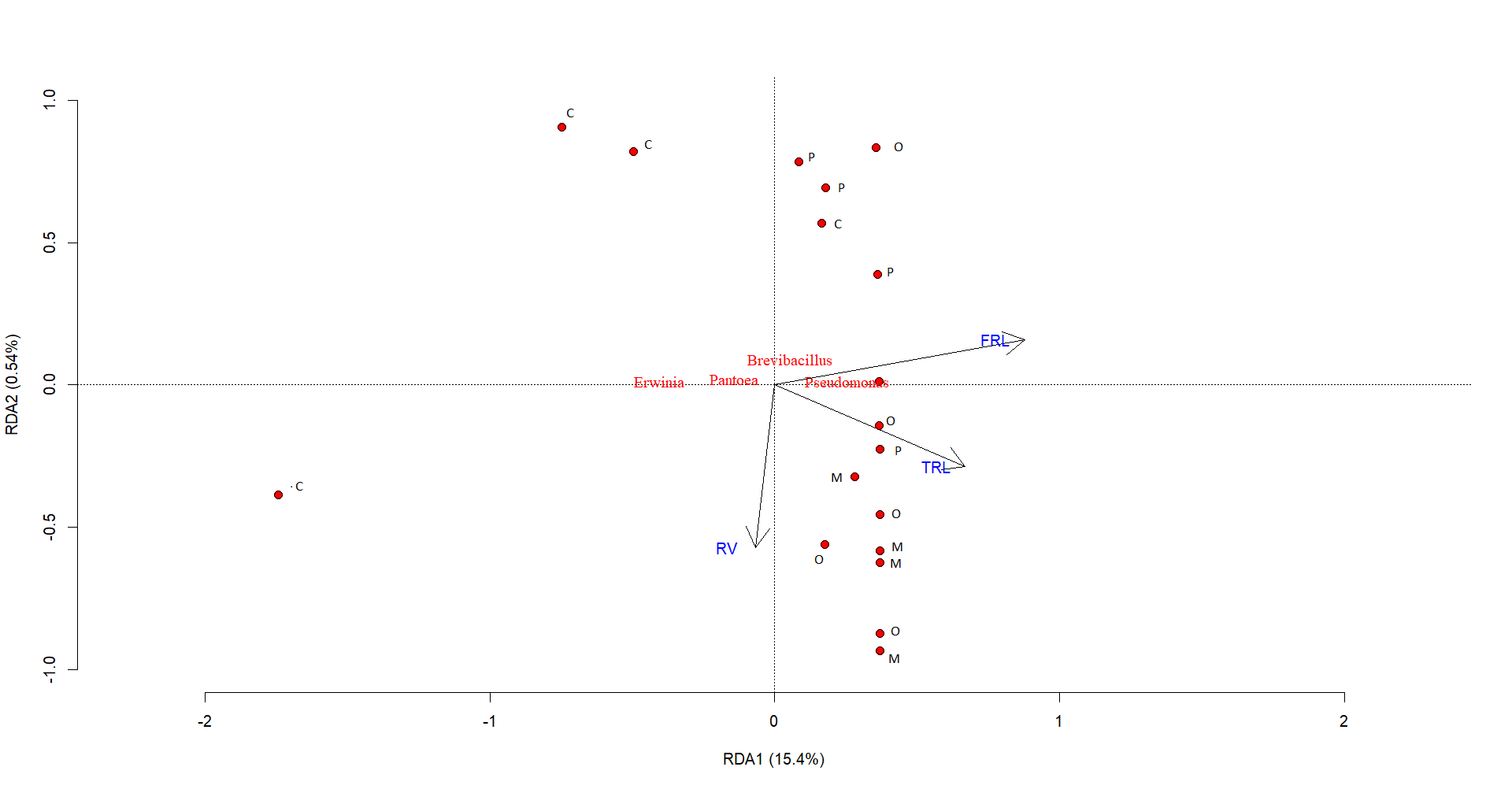
