## Supplementary material for "Genotype-Specific Root Morphology and Metabolic Traits Shape Bacterial Communities and Tolerance to *Fusarium* Root Rot in Wheat": Table S1: Supplementary Table 1.docx

**Supplementary table 1.** Analysis of root dry mass and root morphological traits in 4 wheat varieties at 3-leaf stage. Values represents the means ± standard error of 5 replicates. Means with the same letter are not significantly different, with NS *P*>0.05, * *P*<0.05, ** *P*<0.01 and *** *P*<0.005

| Root traits |  | | Wheat varieties | | | |
| --- | --- | --- | --- | --- | --- | --- |
|  | Concret | Mutic | | Oregrain | Pilier | P-value |
| TRL (cm) | 1455.6±63.7^b^ | 1521.1±110.7^ab^ | | 1855.6±248.27^a^ | 1382.0±168.7^b^ | *** |
| RAD (mm) | 0.41±0.01^a^ | 0.40±0.03^a^ | | 0.38±0.01^a^ | 0.30±0.01^a^ | NS |
| RV (cm^3^) | 1.99±0.14^ab^ | 1.74±0.30^ab^ | | 2.24±0.10^a^ | 1.39±0.10^b^ | * |
| RDW (mg) | 312.0±14.7^a^ | 338.5±31.1^a^ | | 352.9±24.6^a^ | 306.6±21.1^a^ | NS |
| SRL (m.g^-1^) | 4.61±0.30^a^ | 4.82±0.70^a^ | | 5.60±0.60^a^ | 4.53±0.70^a^ | NS |
| RD (g.cm^-3^) | 0.15±0.01^a^ | 0.15±0.04^a^ | | 0.17±0.012^a^ | 0.22±0.04^a^ | NS |
| FRL (cm) | 591.1±65.2^b^ | 687.7±92.4^ab^ | | 821.1±149.7^ab^ | 629.8±43.0^a^ | * |
| CRL (cm) | 894.9±36.3^ab^ | 887.9±76.4^ab^ | | 1033.5±98.8^a^ | 710.3±68.4^b^ | ** |
| FRV (cm^3^) | 0.10±0.00^b^ | 0.11±0.01^ab^ | | 0.15±0.03^a^ | 0.12±0.00^b^ | * |
| CRV (cm^3^) | 2.55±0.20^ab^ | 2.17±0.50^ab^ | | 2.97±0.20^a^ | 1.76±0.20^b^ | * |
| FRR Index | *1.3*±0.42^b^ | *1.6*±0.55^ab^ | | *1.7*±0.41^ab^ | *2.3*±0.41^a^ | **** |
